# Axon-seq decodes the motor axon transcriptome and its modulation in response to ALS

**DOI:** 10.1101/321596

**Authors:** Jik Nijssen, Julio Aguila Benitez, Rein Hoogstraaten, Nigel Kee, Eva Hedlund

## Abstract

Spinal motor axons traverse large distances to innervate target muscles, thus requiring local control of cellular events for proper functioning. To interrogate axon-specific processes we developed Axon-seq, a refined method incorporating microfluidics, RNA-seq and bioinformatic-QC. We show that the axonal transcriptome is distinct from somas and contains fewer genes. We identified 3,500-5,000 transcripts in mouse and human stem cell-derived spinal motor axons, most of which are required for oxidative energy production and ribogenesis. Axons contained transcription factor mRNAs, e.g. Ybx1, with implications for local functions. As motor axons degenerate in amyotrophic lateral sclerosis (ALS), we investigated their response to the SOD1^G93A^ mutation, identifying 121 ALS-dysregulated transcripts. Several of these are implicated in axonal function, including Nrp1, Dbn1 and Nek1, a known ALS-causing gene. In conclusion, Axon-seq provides an improved method for RNA-seq of axons, increasing our understanding of peripheral axon biology and identifying novel therapeutic targets in motor neuron disease.

## INTRODUCTION

Spinal motor neurons (MNs) are highly polarized cells. Their somas and associated dendrites are located in the spinal cord, while their axons traverse the body and connect to distal muscle fibers. The large distance between MN somas and their pre-synapse implies that the distal axon must contain a micro-environment able to independently respond to internal and external triggers. Fast-anterograde transport in axons carries vesicles with transmembrane proteins at rates up to 400 mm per day. However, vesicles containing proteins and RNAs travel considerably slower, at 0.1-10 mm per day (Lasek et al., 1984). Thus, protein transport alone would not suffice to sustain the dynamics of an axon and synapse, implying a strong role for local regulation. Indeed, it is well established that local RNA translation in the synapse is important for temporal control of protein synthesis and synaptic plasticity (Holt and Schuman, 2013).

Motor axons and their specialized synapses with muscle, termed neuromuscular junctions (NMJs), are primary targets in the lethal disease amyotrophic lateral sclerosis (ALS). MNs undergo progressive degeneration in ALS, resulting in denervation of muscle, paralysis and ultimately death at 3-5 years post-diagnosis (Swinnen and Robberecht, 2014). Notably, MNs follow a distinct ‘dying-back’ pattern of degeneration in ALS. Muscle denervation and axonal retraction occur before MN somas in the spinal cord are lost, implying that the NMJ is a highly vulnerable entity of the MN (Comley et al., 2016; Fischer et al., 2004; Valdez et al., 2012). Mutations in several genes such as *SOD1*, *C9orf72*, *TARDBP* (TDP43) and *FUS* have been discovered to cause ALS (DeJesus-Hernandez et al., 2011; Kwiatkowski et al., 2009; Renton et al., 2011; Rosen et al., 1993; Sreedharan et al., 2008; Vance et al., 2009). However, expression of these genes is not restricted to MNs. Why and how they selectively affect MNs in ALS is not clear. Therefore, studying the RNA composition of the distal axon of this vulnerable neuron subtype may illuminate mechanisms driving the specific dying-back degeneration observed in ALS.

Previous efforts to examine axons in isolation *in vitro* have used either cell culture insets, Campenot chambers or microfluidic devices (Boyden, 1962; Campenot, 1977; Taylor et al., 2005). While these are excellent tools for separating axons from somas, residual cross-contamination between the compartments still occurs. In this way, RNA-sequencing efforts aimed at isolating an axonal transcriptome profile (Briese et al., 2016; Gumy et al., 2011; Minis et al., 2014; Rotem et al., 2017; Saal et al., 2014; Taylor et al., 2009) can be easily undermined by a failure to carefully examine the purity of axonal fractions. Due to unappreciated technical difficulties, which are not always acknowledged, axonal fractions can be contaminated with other cellular components or non-neuronal cells. To accurately investigate motor axon mRNA composition and its modulation in ALS we developed Axon-seq, a novel application of our single-cell RNA-preparation technique (Nichterwitz et al., 2016) to microfluidic devices housing mESC derived MNs. In contrast to previous methodologies, Axon-seq does not require RNA isolation, and allows high sensitivity, cost-efficient sequencing from a single microfluidic device. Importantly, Axon-seq utilizes a stringent and sensitive bioinformatic quality control step that identifies samples containing trace levels of mRNA from undesired cell somas, effectively eliminating all cellular cross-contamination.

We derive wild type or ALS (mutated SOD1^G93A^) MNs from mouse embryonic stem cells (mESCs) harboring the MN specific reporter Hb9::eGFP. Using Axon-seq, we show that a pure axonal transcriptome consists of roughly 5,000 mRNA species, compared to >15,000 in cell somas. Approximately 10% of the axonal transcriptome was enriched in the axonal versus the somatodendritic compartment, including transcripts involved in energy production, translation and distinct transcription factor mRNAs such as Ybx1. We also reveal 1,750 mRNAs common to peripheral axons by comparing axonal transcriptome data sets across central and peripheral neuron types. This analysis further decoded a motor axon specific set of 396 mRNAs. Application of Axon-seq to ALS axons uncovered 121 dysregulated transcripts compared to controls, of which several are important for proper neuronal and axonal functioning. Finally, a cross-comparison with axons after *Smn* knockdown, a model for the motor neuron disease spinal muscular atrophy (SMA), revealed that axon guidance receptor neuropilin 1, was downregulated in axons across both MN diseases.

## RESULTS

### Axon-seq is a refined method that enables RNA sequencing of axons from a single microfluidic device

To investigate transcriptional changes in motor axons in health and ALS we used mESC-derived MNs from a control HB9::eGFP line or cells overexpressing the human mutated SOD1^G93A^ protein. We first confirmed the MN subtype identity of our cultures through bulk RNA sequencing and cross-comparison with single MNs (Nichterwitz et al., 2016). We observed expression of the MN transcription factors Isl1 and Foxp1 in our cultures, similar to single cell data on MNs (Supplemental Fig. S1A). This indicates that MNs of a lateral motor column (LMC) identity, which are patterned by Hox genes, were generated (Dasen et al., 2008). Consistent with this, mRNAs of multiple members of the of Hox1-8 families, which define a cervical to thoracic identity, were detected. Further, expression of the transcription factor Phox2b indicates that a proportion of brainstem MNs are also generated (Supplemental Fig. S1A). After plating cells in microfluidic devices, MN axons were recruited to the vacant chamber by a gradient of GDNF and BDNF (Fig. 1A, F). An initial concentration of 50 ng/ml of GDNF/BDNF was followed by a lower concentration of 5ng/ml once axons had crossed the micro-channels, as this avoided growth cone collapse (Fig. 1A). The HB9::eGFP reporter signal was detectable in MN axons, as well as in somas where it co-localized with the MN marker Islet1/2 (Fig. 1B). We next stained for *Map2* (Fig. 1C, Supplemental Fig. S1B) and *Mapt* (Tau) (Fig. 1D, Supplemental Fig. S1B). In mature neurons, *Map2* labels dendrites, while *Tau* labels axons (Garner et al., 1988; Litman et al., 1993). The majority of processes extending across the channels were *Tau*-positive, while a minority were *Map2-positive*. Thus, axonal processes were enriched in this compartment compared to dendrites, likely due to the longer length of motor axons and the relatively long distance of the grooves (Fig 1C-D). MN cultures derived from mESCs also contain interneurons. However, since 90.75% ± 4.8% of Tau^+^ processes were Hb9-GFP+ and only a very low proportion of GFP-negative (and *Tau*-positive) axons appeared in the axonal compartment, we here-on refer to axonal preparations as motor axons.

**Fig. 1.**
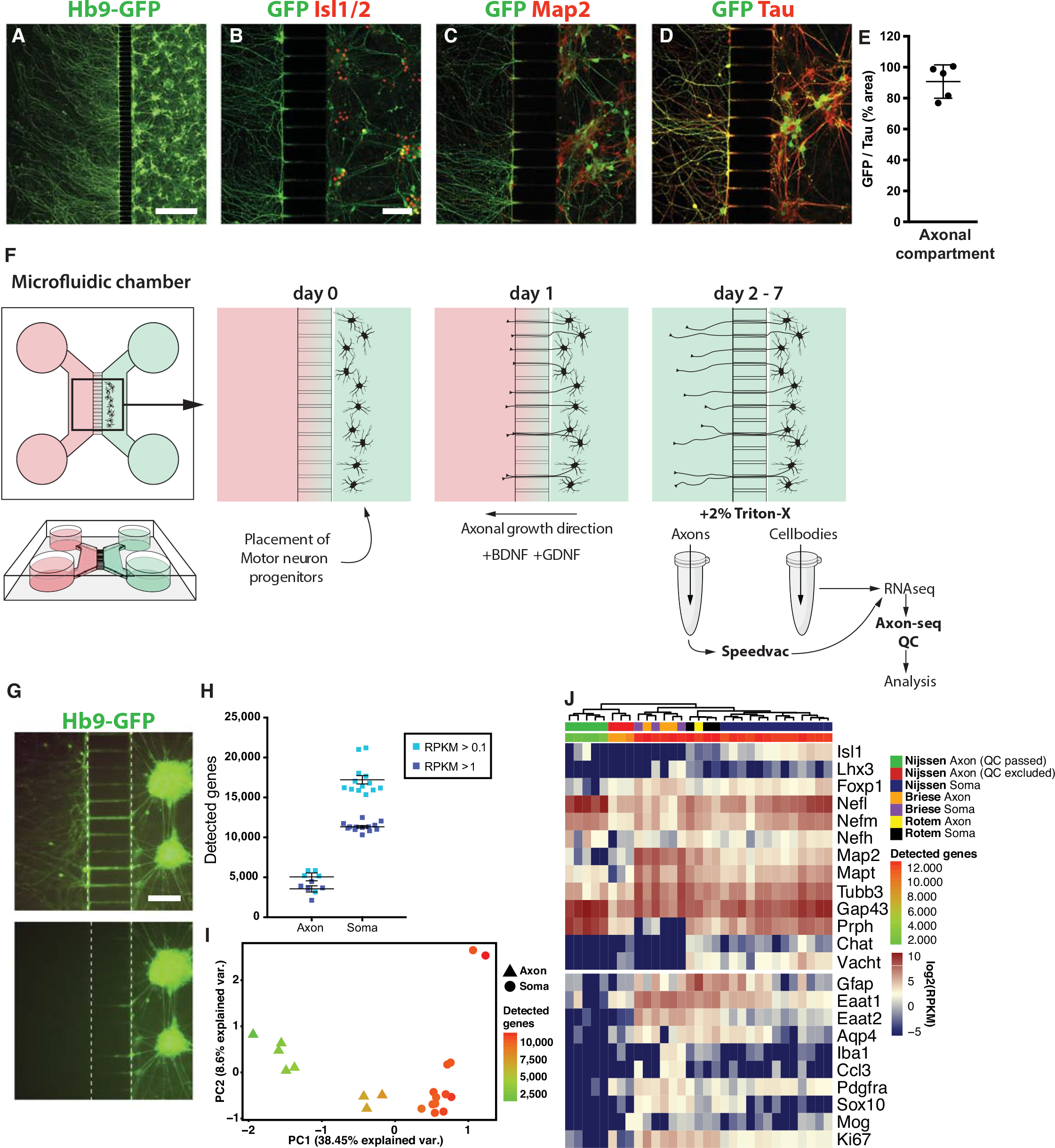
Axon-seq of motor axons in single microfluidic devices. **(A, B)** Anti-GFP immunofluorescence visualizes motor axons crossing the microgrooves and extending into the axonal compartment. MN somas are visualized with Hb9::eGFP and Isl1/2 staining. **(C, D)** Presence of Tau (Mapt) but very little Map2 indicates that mainly axons and not dendrites cross over into the other compartment. Quantification of GFP+ area over Tau^+^ area is shown in **(D)**, revealing that the majority of crossing axons are Hb9::eGFP positive. **(E)** Quantification of the percentage of all Tau^+^ axons that were Hb9::eGFP+ motor axons (90.75% ± 4.8%). Data is represented as mean ± SEM. **(F)** Overview of the used methodology. MNs were cultured in one compartment of the microfluidic device and axons were recruited across microgrooves using a gradient of trophic factors. Compartments were separately lysed and cDNA libraries prepared for RNA sequencing. **(G)** Hb9::eGFP expression reveals that the somatic chamber is not affected by lysis of the axons. **(H)** The axonal fraction contained around 5,000 unique transcripts, while MN somas contained >15,0 (mean ± SEM). **(I)** PCA based on all expressed genes showing ‘contaminated’ axon samples with an intermediate amount of detected genes that cluster away from the clean axon samples and towards the somatodendritic samples. **(J)** Expression of selected marker genes in soma and axon samples of this study and data from studies by Rotem et al. (2017) and Briese et al. (2016). Our clean axon samples separate from the rest, while axon and soma samples from previously published studies intermingled, indicating crosscontamination. Scale bars: **A:** 500 μm. **B:** 100 μm, also applies to **C** and **D**. **G:** 100 μm.

Adding different media volumes to the microfluidic compartments can generate a fluid flow that counteracts diffusion (Mills et al., 2018; Taylor et al., 2005). Using this principle, axonal or somatodendritic samples were lysed without affecting the opposite compartment (Fig. 1G). Further, both compartments could be effectively lysed using 2% triton X-100 in water (Fig. 1F, G), eliminating the need for RNA purification. To boost final RNA concentration and thereby allow sequencing from an individual microfluidic device, we introduced a vacuum centrifugation concentration step, allowing the entire lysate to be used for subsequent cDNA library preparation. Using this cost-efficient approach, we successfully conducted polyA-based RNA-seq on axonal and soma samples from individual devices using the Smart-seq2 protocol (Picelli et al., 2013) with some minor modifications (Nichterwitz et al., 2016).

We next implemented a crucial bioinformatics-based quality control step (Axon-seq QC) to exclude axon samples displaying trace levels of cross-contamination from the adjacent compartment. First, to be considered for further analysis, samples were required to display at least 300,000 mapped reads, a minimum of 0.4 correlation to at least one other sample, and more than 2,500 detected genes (See Methods). Secondly, we performed principal component analysis (PCA) based on all expressed genes. In this analysis PC1 strongly reflected the number of detected genes, where axonal transcriptomes occupied the lower ranges, and somatodendritic samples occupied the higher ranges (Figure 1I). Three axonal samples displayed 7,539, 8,485 and 7,760 detected genes and occupied mid PC1 ranges. These samples were likely contaminated with one or more soma and were discarded. Remaining high quality axon samples contained roughly 3,500 detected genes at > 1 RPKM and 5,000 detected genes at > 0.1 RPKM, while somatodendritic samples contained approximately three times these numbers (Fig. 1H). Axonal samples also displayed fewer detected genes when compared to single MNs (Nichterwitz et al., 2016) (Supplemental Fig. S1F). Interestingly, despite this difference, cDNA yield was comparable between axonal samples and single MNs (Supplemental Fig. S1E), suggesting that *i)* the cDNA yield from all axons in a microfluidic device approaches that of at most only a few cells, and *ii)* low numbers of detected genes in these samples is not a technical artefact, and is instead a trait unique to the axonal transcriptome. Given that Axon-seq libraries display lower numbers of detected genes than do single cells, a single contaminating cell has potential to drastically and unacceptably alter the readout of an axonal library. We find that consideration of the number of detected genes is perfectly suited to identifying these contaminated samples, allowing further downstream analysis on only pure Axon-seq samples.

After QC, we then sought to compare our Axon-seq dataset to previously published methods, selecting two RNA-seq studies (Supplemental Table S2) that used either cell culture insets (Rotem et al., 2017) or single microfluidic devices (Briese et al., 2016) to isolate motor axon transcriptomes. Both these studies display good quality data and describe enrichment of axonal transcript. Expression of neuronal markers in all samples was visualized in a heatmap, including e.g. neurofilaments, peripherin, *Mapt*, *Gap43* and the glial markers *Gfap*, *Aqp4*, *Eaat1*, *Iba1*, *Ccl3*, *Pdgfra*, *Sox10* and *Mog* (Fig. 1J). This analysis clearly demonstrates that our Axon-seq samples that passed the QC expressed robust levels of neuronal and axonal markers, while being devoid of glial markers (e.g. Gfap, Eaat1, Pdgfra and Sox10) and of the proliferative marker Ki67, which were in fact present in all previously published axon sequencing samples (Fig. 1J). Furthermore, our axon samples clustered away from somatodendritic and our cross-contaminated axon samples, while axon and soma samples from previously published studies were intermingled (Fig. 1J). Further cross-comparison of our axon samples with a published data derived from purified neuronal samples (Zhang et al., 2014) confirmed their purity (Supplemental Fig. 1D). Importantly, our Axon-seq samples expectedly displayed a far lower number of detected genes than the previously published axon samples (Fig. 1J, top color bar). Thus, samples displaying even small levels of soma contamination were successfully identified through application of Axon-seq QC.

In summary, our analysis demonstrates the importance of extensive QC of axon transcriptome data with particular consideration for the number of detected genes and presents Axon-seq as a robust and significantly improved method for RNA-seq of pure axonal fractions from individual microfluidic devices and sensitivity similar to single cell sequencing.

### Axons have a unique transcriptional profile and are particularly enriched in transcripts important for mitochondrial and ribosomal functions

We next sought to compare the transcriptome of the axonal fraction against the somatodendritic fraction. When considering transcripts present in at least 3 out of 5 axonal samples, the axonal transcriptome comprised 4,238 genes (Supplemental Table S3). 10,406 genes were differentially expressed between axons and somas. The majority of these (9,762) were enriched in the soma, while 644 transcripts were axon-enriched (Supplemental Table S4). Moreover, analysis of the top 100 differentially expressed genes in axons and somas respectively, clearly demonstrated that axons are not simply diluted somas, but instead have a unique mRNA profile (Supplemental Fig S1G, H). Principle component analysis based on all expressed genes clearly distinguished axons from somas, with PC1 explaining 41.7% of the variance (Fig. 2A). The top 20 genes determining PC1-negative loading and thereby axonal identity, included several genes involved in the mitochondrial respiratory chain (*Cox6a1*, *Cox6b1*, *Cox8a*, *Cox4i1*, *Ndufa3*, *Ndufa7*, *Ndufb9*, *Ndufb11*, *Uqcrq*), others in mitochondrial ATP synthesis (*Atp5k, Atp5j2*), ribosome subunits and transcriptional elongation (*Rpl41*, *Tceb2*), cytoskeleton organization (*Tmsb10*), copper delivery (*Atox1*) and mRNA splicing (*Ybx1*) (Fig. 2B). The PC1-positive loading (Fig. 2A) was strongly influenced by transcripts present in somas that were completely absent in axons. In total, 2,903 genes were expressed at an average of RPKM > 1 in somas, that were undetectable in axons (Supplemental Fig S1H). Gene set enrichment analysis (GSEA) on axonal enriched genes uncovered pathways related to both local translation (Fig. 2C, D), mitochondrial oxidative energy production (Fig. 2E) and nonsense-mediated decay (Fig. 2F). Interestingly, both nuclear- or mitochondrial encoded transcripts were enriched in axons (Fig. 2G, H) but the ten most abundant energy production-related transcripts in axons were nuclear encoded transcripts (Fig. 2I).

**Fig. 2.**
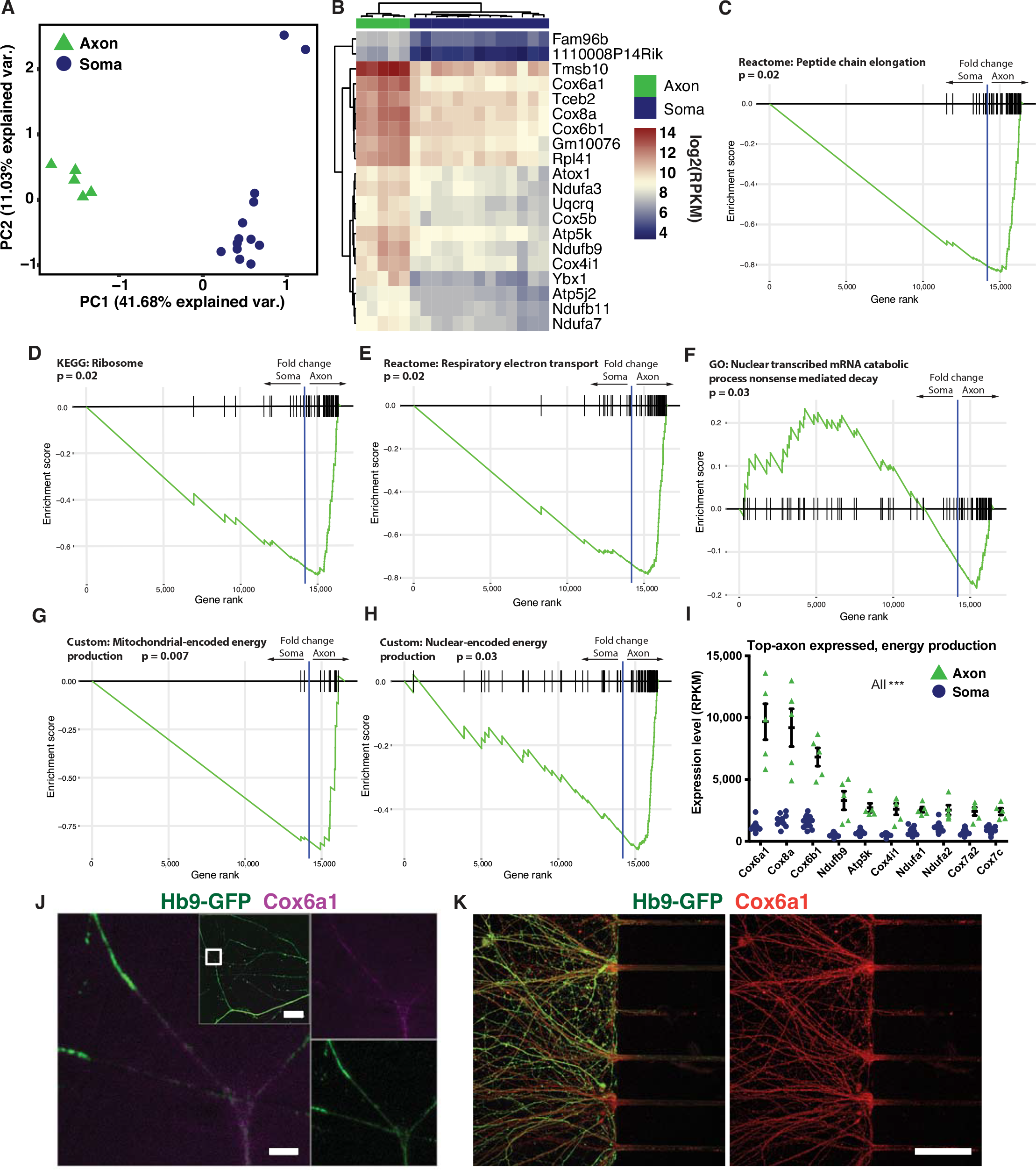
Axons have a distinct transcriptome from somas. **(A)** PCA of all expressed genes revealed a clear separation of axons and somas. **(B)** Top 20 genes determining PC1-negative loading. All of them are strongly enriched in axons compared to somas. (**C-H)** Gene set enrichment analysis (GSEA) using fold changes from the DESeq2 differential expression output ranked list between axons and somas, without a p-value cut-off. GSEA uncovered pathways related to local translation **(C, D)**, energy production **(E)** and nonsense-mediated decay **(F)**. Subdividing the oxidative energy production class into mitochondrial- and nuclear-encoded transcripts reveals an axonal enrichment for both **(G, H)**. Differential expression was performed on the 5 axon samples that passed QC and 13 soma samples, as visualized in panels **A** and **B**. **(I)** Expression levels of the highest expressed transcripts in axons related to oxidative energy production. All of the genes plotted are significantly enriched in the axonal compartment compared to the somatodendritic compartment. P-values are derived from the DESeq2 differential expression and were adjusted for multiple testing. Data is represented as mean ± SEM. **(J, K)** RNAscope **(J)** and immunocytochemistry **(K)** of Cox6a1, the most abundant mRNA in axons involved in energy production. Scale bars: **J:** 2 μm, lower magnification insert: 20 μm. **K:** 50 μm.

To confirm the axonal localization of particular transcripts we conducted *in situ* hybridization experiments using RNAscope. Control experiments using negative (DapB) and positive (Ubc, Polr2a and Ppib) control probes demonstrated the specificity of the method (Supplemental Fig. S2A, B). Using an oligo dT probe we could show that poly-A mRNAs were present in axons (Supplemental Fig. S2C). In subsequent experiments, we confirmed that Cox6a1 mRNA, which is part of the electron transport chain, was localized to Hb9-GFP+ motor axons (Fig. 2J), as well as to somas (Supplemental Fig. S2D). Immunocytochemistry of motor axons demonstrated that also the *Cox6a1* protein was readily detected in axons (Fig. 2K).

In summary, our data shows that the axons of growing MNs are enriched for mRNAs related to ribosomes and mitochondrial oxidative phosphorylation, a large majority of which are mitochondrially produced. The distal axon compartment can thereby sustain local translation machinery and high energy metabolism, reflecting the enormous energy demands of growing axons.

### Motor axons show a unique transcription factor profile

Analysis of the top 50 most prevalent transcription factor mRNAs in each of the two compartments uncovered 16 common transcription factors, including *Ybx1*, *Carhsp1* and *Sub1* (Fig. 3A, B). Notably, the majority of soma-enriched TFs mRNAs were more abundant in somas than most axon-enriched TFs were abundant in axons (Fig. 3B). However, five of the ten most abundant axonal TF mRNAs were significantly enriched in this compartment compared to somas, including *Ybx1, Carhsp1, Pbx4, Hmga1* and *Zfp580* (Fig. 3C). The enrichment of *Ybx1* (Y-box binding protein 1, *p* < 0.001) (Fig. 3C) was particularly compelling since Ybx1 partakes in e.g. pre-mRNA transcription and splicing, mRNA packaging and regulation of mRNA stability and translation (Lyabin et al., 2014a; Lyabin et al., 2014b). RNAscope confirmed the presence of Ybx1 mRNA in Hb9-GFP+ motor axons (Fig. 3D). Immunocytochemistry demonstrated that also the Ybx1 protein was localized to motor axons (Fig. 3E), strengthening a possible role for Ybx1 in supporting local regulation of distal axonal cellular processes.

**Fig. 3.**
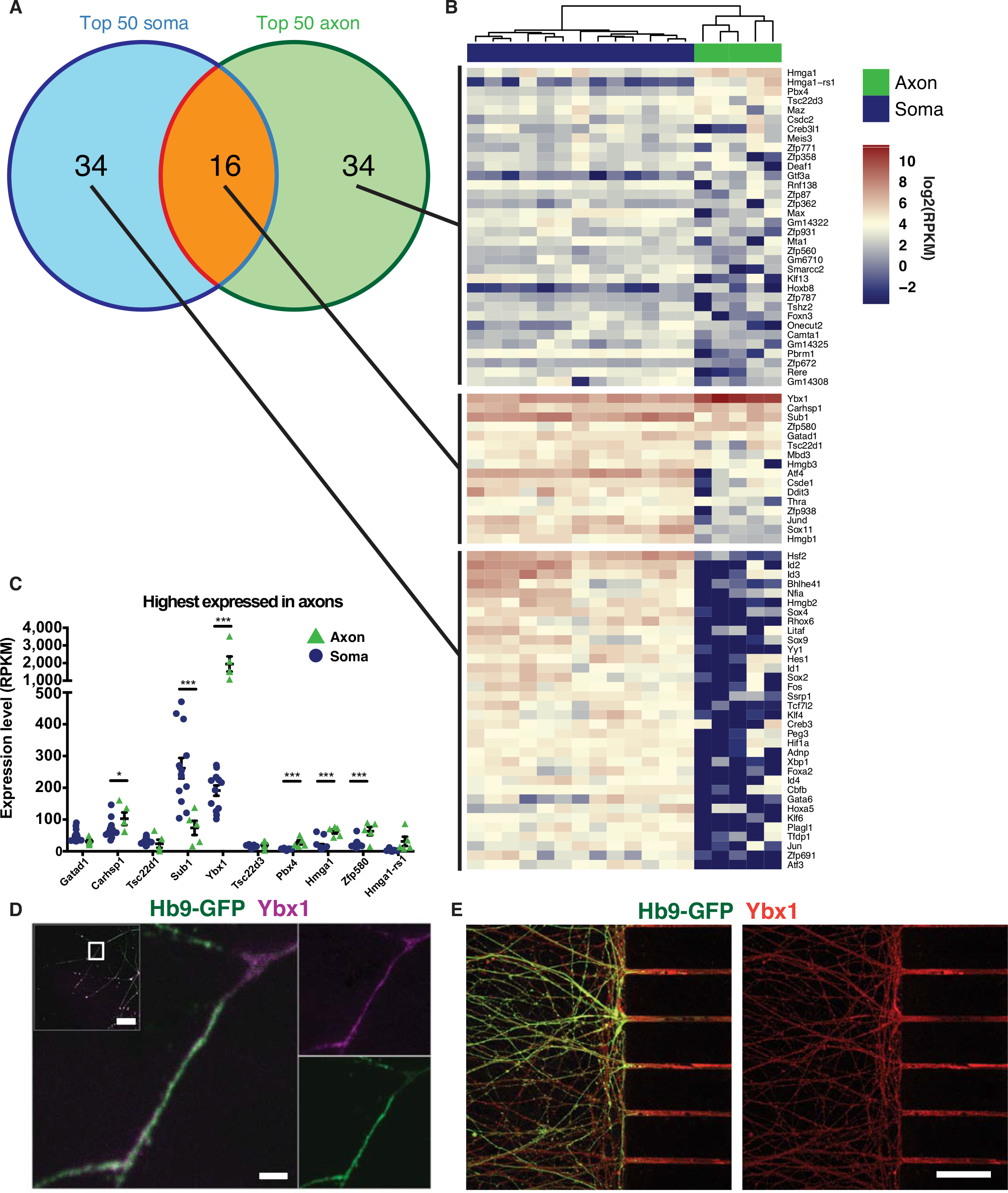
A unique transcription factor repertoire localizes to axons. **(A)** Of the top 50 highest expressed transcription factors in the soma and axon (at least 4 axonal samples), 16 overlapped. **(B)** Heatmap of expression values of transcription factors from panel A. **(C)** Top 10 transcription factors with the highest expression in axons. 5 were significantly enriched in axons, while 1 was enriched in somas. P-values are derived from differential expression between the 5 axon and 13 soma samples that passed QC and are corrected for multiple testing (mean ± SEM). **(D, E)** RNAscope FISH **(D)** and immunocytochemistry **(E)** for Ybx1, the most abundant transcription factor at mRNA level in axons. Scale bars: **D:** 2 μm, lower magnification insert: 20 μm. **E:** 50 μm.

In summary, axons show a unique transcription factor profile which is expected to reflect functions particular to this compartment, and that could potentially function as communication and control signals between somas and their long range axonal projections.

### Cross comparison of axonal data sets across neuronal subtypes reveals both motor neuron-enriched and pan-neuronal axon mRNA repertoires

To understand which mRNAs are specific to motor axons versus transcripts defining growing axons in general, we compared our Axon-seq motor axons with a published axon dataset derived from primary embryonic mouse MNs (Briese et al. 2016), and with primary embryonic mouse dorsal root ganglia (DRG) (Minis et al., 2014). In both comparisons, GO-term analysis on the overlapping genes reveals oxidative energy production, translation and localization of proteins to mitochondria and ribosomes. (Supplemental Fig. S3A and B, Supplemental Table S6 and S8). Both the primary motor axon and primary DRG axon data sets contained a large number of proliferative and glial marker sets that were absent from our Axon-seq data (Fig. 1J, Supplemental Fig. S3C). This was further confirmed by GO term analysis of the transcriptomes unique to *Briese et al.* and *Minis et al.*, which in both cases identified biological processes involved in cell division (Supplemental Fig. S3A and B, Supplemental Table S7 and S9).

Notably, Cox6a1 and Ybx1, abundant mRNAs in motor axons were present across neuron types. Furthermore, a lowly expressed, but important transcript, Nrp1, was also pan-neuronal (Supplemental Fig. S3A and B). We could confirm the presence of this transcript in our motor axons using RNAscope (Supplemental Fig. S3D).

In summary, our analyses have identified a general axonal transcriptional signature as well as defined a unique motor axon code which gives clues to function as well as their particular susceptibility in disease.

### Axon-seq of human motor axons identifies a common transcriptome with mouse motor axons

We further applied Axon-seq to human MNs derived from two control induced pluripotent stem cell (iPSC) lines and grown in microfluidic devices. Human motor axons could be readily recruited across microgrooves using a BDNF/GDNF gradient (Fig. 4A). In a PCA plot, the axonal samples clearly separated from human somatodendritic fractions (Fig. 4B). Considering transcripts present in at least three axonal samples, the human motor axonal transcriptome was comprised of 2,793 genes (Supplemental Table S10). The number of detected genes was on average 2,611 ± 286 at RPKM > 1, and 3,495 ± 460 at RPKM > 0.1 (Fig. 4C). Human motor axons were strongly enriched for cytoskeletal transcripts such as NEFL, NEFM and GAP43 and showed negligible glial contamination was present (Fig. 4D).

**Fig. 4.**
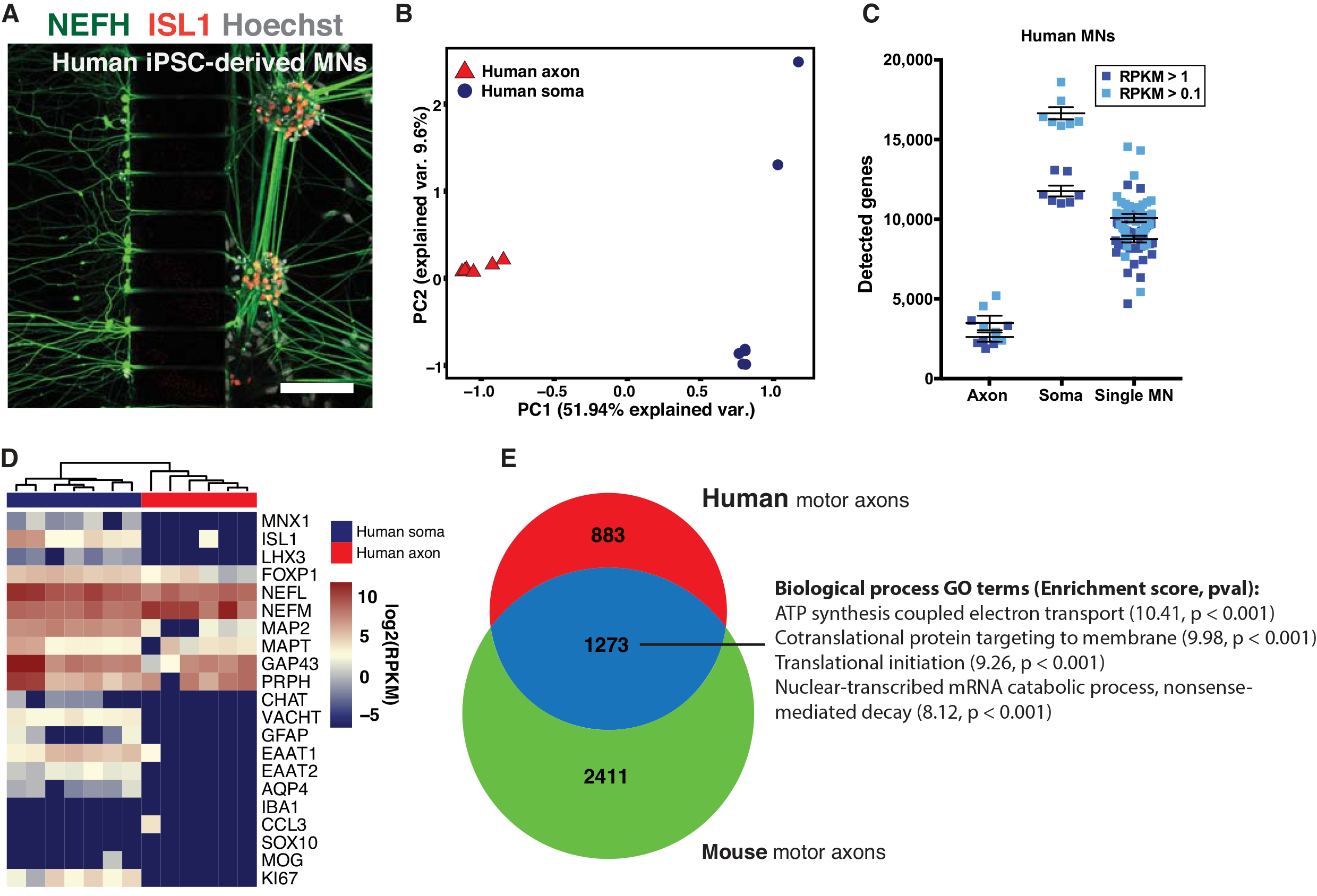
Human motor axons have a distinct transcriptome that overlaps with mouse motor axons. **(A)** Immunocytochemistry of human control iPSC-derived motor neurons growing in a microfluidic device. **(B)** PCA based on all expressed genes shows clear separation of human axonal and somatodendritic fractions along the first principal component. **(C)** Numbers of detected genes in human bulk somas and axons, as well as single human Hb9::GFP+ motor neurons. Human axons contain > 1,000 transcripts, while somas contain > 15,000 transcripts at RPKM > 0.1. Single cells are intermediate with approximately 10,000 transcripts detected. **(D)** Expression of selected marker genes in human soma and axon samples reveals strong axonal presence of mRNAs for cytoskeletal elements, as well as the transcription factor FOXP1. Glial markers were absent in axons. **(E)** Comparison of the human motor axon transcriptome with the common mouse motor axon transcriptome (Supplemental Fig. S3A) reveals an overlap of 1273 genes (after exclusion of non-orthologous genes). These genes enrich in GO terms for energy production, translation and nonsense-mediated decay. Scale bars: **A:** 100 μm.

We sought to investigate the overlap between our human motor axon transcriptome and the previously defined common mouse motor axon transcriptome (Supplemental Fig. S3A). Selecting only the genes present in three or more samples for each species and excluding any genes that did not have annotated orthologs in human and mouse, we found 1,273 genes to be common between human and mouse motor axons. GO-term analysis of these genes revealed energy production, translation and nonsense-mediated decay as enriched processes (Fig. 4E, Supplemental Table S11).

### The ALS-causative mutation SOD1^G93A^ modulates the axonal transcriptome

Motor axons are early pathological targets in the lethal MN disease ALS, where die-back pathology begins in the distal axon. We therefore investigated if overexpression of an ALS causative mutation (*SOD1*^*G93A*^) would induce changes in the axonal transcriptome. MNs derived from mESCs overexpressing human SOD1^G93A^ have high levels of SOD1 protein in somas and axons (Fig. 5A). Axon-seq of SOD1^G93A^ axons identified 4,479.5 ± 1327.7 genes at RPKM > 1 and 6,568.8 ± 2293.1 at RPKM > 0.1 (Supplemental table S12). Human mutant *SOD1* was highly expressed in somas and axons (Fig. 5B) at levels approximately 10-fold higher than mouse *Sod1* (Fig. 5C). Interestingly, mouse *Sod1* mRNA levels were higher in axons than in soma (p < 0.001), likely reflecting the important role of this enzyme in converting reactive superoxide radicals produced in the abundant axonal mitochondria. Mutant *SOD1* overexpression resulted in differential expression of 121 genes in axons compared to the control line, of which 96 were upregulated in SOD1^G93A^ axons, while 25 were reduced (Fig. 5D, Supplemental Table S13). Strikingly, all but two of these genes (*Zfand1* and *Zfp688*) were uniquely dysregulated in mutant axons and were not differentially expressed between the control and SOD1^G93A^ soma compartments (Supplemental Table S14). Moreover, many of these genes have previously been implicated in general neuronal function as well as in ALS pathology (Fig. 5E). For instance, we found that several transcripts paramount for proper axon function were downregulated or completely absent in SOD1^G93A^ axons, including *Dbn1* (Lin and Koleske, 2010; McIntyre et al., 2010; Sharma et al., 2012), *Nrp1* (neuropilin 1) (Telley et al., 2016) and *Mgrn1* (Upadhyay et al., 2016). Some transcripts that were upregulated in SOD1^G93A^ axons appear detrimental, including *Adgr1* (Zuko et al., 2016) (Fig. 5E). Additionally, upregulation of some transcripts in ALS could reflect compensatory mechanisms, including *Dhx36* (Bicker et al., 2013), the ALS-causative gene *Nek1* (Kenna et al., 2016), *F3* (contactin 1) (Bizzoca et al., 2009), *Rbpms* (Hornberg et al., 2013) and *Farp1* (Zhuang et al., 2009) (Fig. 5E), that may prevent axon damage or dysfunction in SOD1^G93A^ axons.

**Fig. 5.**
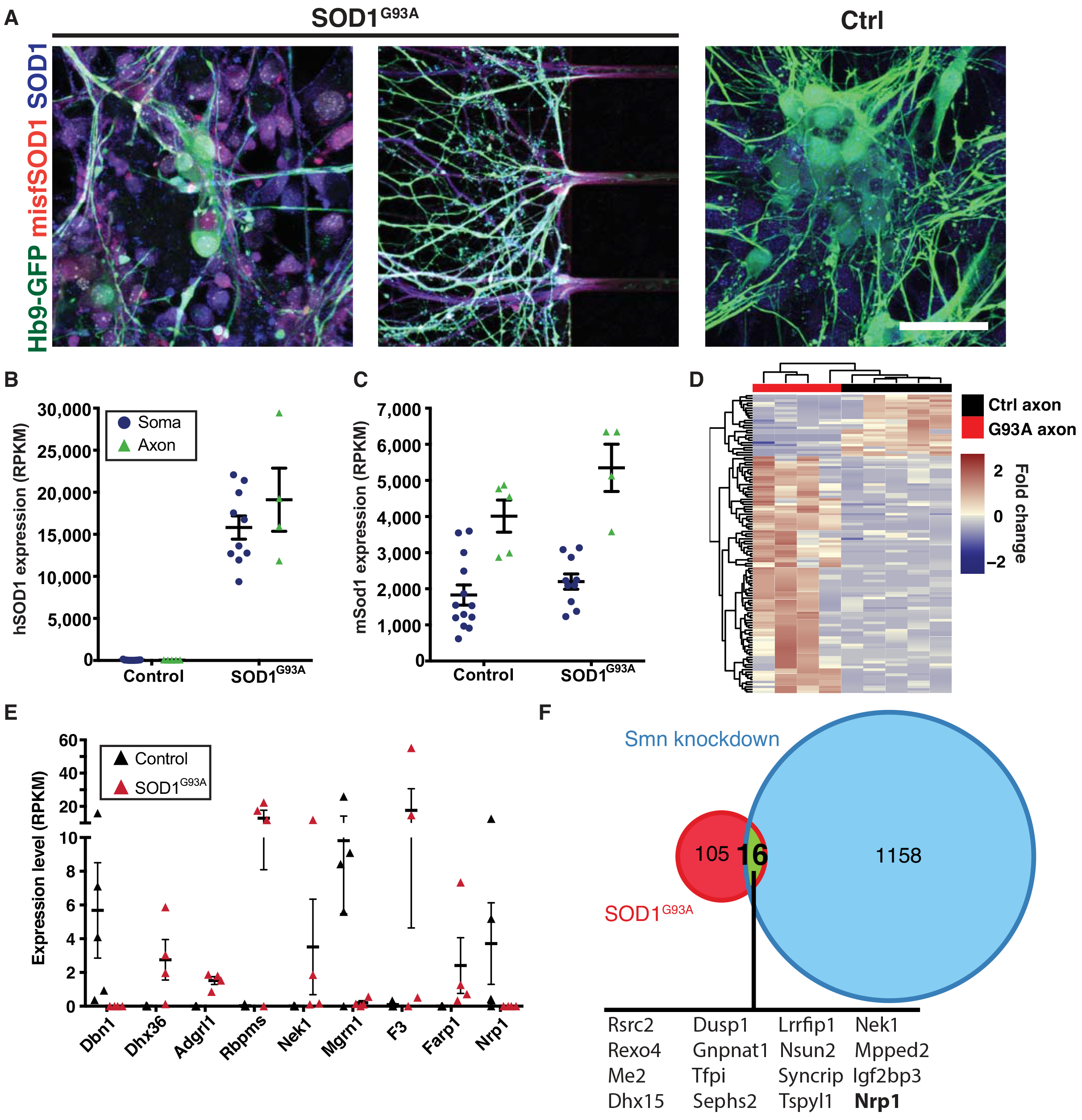
Overexpression of SOD1^G93A^ induces changes in the axonal transcriptome. **(A)** Staining for SOD1 and misfolded SOD1 in MN cultures. In SOD1^G93A^ expressing cells high levels of both SOD1 and misfSOD1 are detected, whereas no misfSOD1 is detected in control cells. **(B, C)** Expression levels of mouse SOD1 and human transgenic SOD1. Mouse SOD1 is enriched in axons, while transgenic human SOD1 is equally divided over both compartments. Levels of hSOD1 are approximately 10-fold higher in somas than mouse SOD1. Data is represented as mean ± SEM. **(D)** Heatmap of differentially expressed genes between 5 control axon samples and 4 axon samples from SOD1^G93A^ overexpressing MNs. A total of 121 genes were differentially expressed, of which 25 enriched in control axons and 96 in SOD1^G93A^ axons. **(E)** Selected differentially expressed genes from the analysis visualized in panel **D** with relevance to ALS and general neuronal functioning. All p-values are derived from the DESeq2 output and are corrected for multiple testing (mean ± SEM). **(F)** Overlap between dysregulated genes upon SOD1^G93A^ expression in our dataset and Smn knockdown in data from Saal et al. (2014). 16 genes are commonly dysregulated. 1 gene, Nrp1, is commonly downregulated.

Finally, we wanted to elucidate if any of the transcripts that were dysregulated by the SOD1^G93A^ mutation were affected across motor neuron diseases. We therefore compared our RNA sequencing data with a microarray data set on motor axons in which the *Smn* gene was knocked down to model the disease spinal muscular atrophy (Saal et al., 2014). Sixteen transcripts were regulated across the two disease models. However, 15 genes were regulated in the opposite direction between the ALS and the SMA disease models, while one transcript, the axon guidance receptor Neuropilin 1 (*Nrp1*), was down-regulated in both motor neuron disease models (Fig. 5F).

In summary, application of Axon-seq methodology to a mutant *SOD1* overexpression model of ALS identified dysregulation of 121 axonal specific transcripts, some of which appear detrimental, while others may indicate compensatory mechanisms to counteract any induced dysfunction. *Nrp1* was downregulated in motor axons across ALS and SMA disease models and might have implications for pathology.

## DISCUSSION

Axonal RNA biology is increasingly implicated in ALS, which recent studies have aimed to address (Briese et al., 2016; Rotem et al., 2017). In this study, we developed Axon-seq, an improved method allowing isolation and interrogation of the transcriptome of distal motor axons. We show that with careful bioinformatic-QC, this cost-effective and sensitive methodology can reliably yield high quality, and pure axonal transcriptomes free of contaminating astrocytic or soma restricted transcripts. Importantly, Axon-seq was successfully applied to both mouse and human motor neurons derived from multiple pluripotent sources and cultured in microfluidic devices. We detected 3,500-5,000 mRNA transcripts in axons, as opposed to somas, where we detected > 15,000 transcripts. Differential gene expression shows that the axonal compartment is not simply a dilution of the soma compartment, and is instead enriched for mRNAs coding for mitochondrial energy production and ribosomes. This distinct profile reflects the high energy-dependence of growing axons and the importance of local protein translation to react to the local environment. For instance, during axon guidance the growth cone responds to guidance cues through local protein synthesis, which is required for directional turning and collapse response (Campbell and Holt, 2001). In fact, axonal protein synthesis in chick sympathetic axons was shown to account for 5% of the total protein synthesis of the cell (Lee and Hollenbeck, 2003). Furthermore, transcripts for all essential subunits of the proteasome system (*Psma1-7* and *Psmb1-7*) were present in motor axons (Supplemental Table S3). Many regulatory subunits and chaperones were also detected, implying extensive regulation of protein degradation in axons. Given that small amounts of RNA can be translated into multiple protein copies, maintaining in neuronal processes an mRNA pool poised for translation both saves energy, and increases capacity for a rapid response. Moreover, some proteins may be toxic during transport and producing these locally would avoid any detrimental effects (Besse and Ephrussi, 2008; Du et al., 2007).

The two largest groups of mRNAs identified in mouse and human axons through GO term analysis and gene set enrichment analysis (GSEA) were mitochondrially-encoded mRNAs involved in energy production and mitoribosomes. As mitochondria are abundant in axons, and assuming there exists no bias in soma to axon transport of mitochondrially-encoded RNAs, we speculate that these mRNAs are locally transcribed. This would be consistent with our finding that nuclear encoded mRNAs for energy production, and in particular nuclear encoded ribosomal RNAs, were less abundant in axons compared to mitochondrially encoded genes, and in agreement with previous reports also describing a predominance of mitochondrially encoded genes in DRG axons (Minis et al., 2014). Mitochondria can replicate by fission in peripheral axons (Amiri and Hollenbeck, 2008), a process that would be facilitated by local transcription and translation of mitochondrially encoded proteins. Interestingly, our data suggests that in fact the complete mRNA library required for mitochondrial replicative fission and function is locally present in axons, in part through a significant contribution from local axonal transcription.

Transcription factors, induced by local morphogens, define the identity of neurons during early development and are later often required for neuronal survival (Kadkhodaei et al., 2013; Sgado et al., 2006). Particular transcription factors have recently been shown to have functions outside of the cell soma and to be important in neuronal circuitry plasticity, axon pathfinding and neuroprotection (Sugiyama et al., 2008; Torero Ibad et al., 2011; Wizenmann et al., 2009). In fact, multiple studies have identified axonal mRNAs for transcription factors, in addition to nuclear transport machinery and the nuclear envelope, whose local translation and subsequent signaling to the soma appear vital to axon maintenance and survival in injury paradigms (Ben-Yaakov et al., 2012; Cox et al., 2008; Hanz et al., 2003; Ji and Jaffrey, 2012; Yoon et al., 2012). Axon-seq detected a specific set of transcription factor mRNAs enriched in distal MN axons compared to somas, including the RNA-binding factor *Ybx1*. YBX1 has known roles in binding and stabilizing cytoplasmic mRNAs as well as in regulating translation by affecting the interaction between eukaryotic initiation factors and mRNAs (Lyabin et al., 2014b). YBX1 was also shown to mediate the anterograde axonal transport of transcripts of the mitochondrial protein, Cox4 (Kar et al., 2017). Interestingly, in this function YBX1 was shown to interact with FUS, an RNA binding protein and known ALS-causing gene (Groen et al., 2013). We expect that YBX1 and other axonally located transcription factors identified here will emerge as important mediators of communication between somas and axons.

‘Dying-back’ pathology in vulnerable MNs imply that the distal axonal compartment undergoes pathological changes during ALS. Axon-seq on motor axons harbouring the ALS-causative mutation SOD1^G93A^ uncovered differential expression of 121 mRNAs, several of which are crucial for neuronal function, axon maintenance and growth. For example, a number of the transcripts that were upregulated in SOD1^G93A^ axons appear detrimental, including; *Adgr1*, which can induce apoptosis and cause reduction in neurite outgrowth (Zuko et al., 2016). Furthermore, multiple transcripts that are important for axon function were downregulated or absent in SOD1^G93A^ axons, including; *Dbn1* which is important for axon initiation, growth and guidance, (Lin and Koleske, 2010; McIntyre et al., 2010; Sharma et al., 2012), *Nrp1* (Neuropilin 1) which is a semaphorin receptor involved in both axon guidance and subcellular target recognition (Telley et al., 2016); and *Mgrn1* a ubiquitin-ligase, which appears important for mitochondrial function and neuronal survival (Upadhyay et al., 2016). *Mgrn1* has previously been shown to be downregulated in SOD1 mice, as well as being recruited to SOD1-positive inclusions (Chhangani et al., 2016). *Nrp1* acts an axonal attractant during development (Chauvet et al., 2007) is important for limb innervation (Huettl et al., 2011). The downregulation of Nrp1 in our study here would thus appear to align with a loss of NMJ function. Nonetheless, it was recently shown that i.p. delivery of an anti-Nrp1 antibody to SOD1^G93A^ mice could lengthen their life-span and reduce NMJ denervation (Venkova et al., 2014). Although it is not evident if down-regulation of Nrp1 within MNs induced the rescue in the *Venkova et al* study, this could that the downregulation we see may be a compensatory event that in some ways could be beneficial, but this remains to be further investigated. It is noteworthy that *Nrp1* was downregulated in motor axons across SMA and ALS disease models. While SMA is generally an early onset motor neuron disease and ALS is a disease correlated with aging, both are characterized by an early motor axon pathology (Cifuentes-Diaz et al., 2002; Comley et al., 2016; Fischer et al., 2004).

Our data combined with published studies indicate that NRP1 might be an early target in ALS pathology and that it potentially could be modulated in both diseases with beneficial results.

Interestingly, the SOD1 mutation also caused an upregulation of transcripts that could be beneficial to axons, possibly through compensatory mechanisms aimed at preventing axon damage or dysfunction that are in play in SOD1^G93A^ axons. Induced transcripts with potential beneficial properties included; *Dhx36*, an ATP-dependent helicase, which mediates dendritic localization of miR-134 with consequent effects on synaptic protein synthesis and plasticity (Bicker et al., 2013); *F3* (contactin 1), a cell adhesion molecule involved in neurite outgrowth (Bizzoca et al., 2009); *Rbpms*, an RNA-binding protein with importance for RNA-granule localization and dendritic arbor complexity, as shown in retinal ganglion cells (Hornberg et al., 2013); *Farp1*, a marker for LMC MNs which positively regulates dendrite growth (Zhuang et al., 2009); *Nek1*, which has a putative role in microtubule stability, neuronal morphology and axon polarity. Interestingly, loss of function variants of *NEK1* confer susceptibility to ALS in humans (Kenna et al., 2016).

In summary, we have developed Axon-seq, a refined method with high sensitivity for RNA sequencing on axons, which contains crucial bioinformatics quality control steps to identify high quality and improved purity axon transcriptomes. Using Axon-seq, we demonstrate that mouse and human motor axons contain a smaller and distinct transcriptome as compared to somas, with high enrichment for transcripts required for local energy production and protein translation. We further show that the majority of these motor axonal genes are shared with axons of other neuronal types, underlining their importance in general axonal biology. Finally, motor axons expressing ALS-causative SOD1^G93A^ displayed both down- and upregulation of key transcripts involved in axonal growth and guidance, as well as neuronal survival. These differences likely represent pathological as well as compensatory processes induced by the mutation.

Due to its robustness, sensitivity and cost effectiveness, Axon-seq is a method of broad utility for the study of RNA dynamics in any polarized cell. Here we successfully applied Axon-seq to human motor axons demonstrating its usefulness for the study of human neuronal processes in stem cell-derived *in vitro* systems. Thus far, axons of human stem cell-derived glutamatergic neurons have been isolated in microfluidic devices and screened using microarrays (Bigler et al., 2017), but not using more advanced RNA-seq methods. Microfluidic devices have also been used to investigate axonal trafficking deficits and axonal degeneration in stem cell-derived MNs from ALS patients (Naumann et al., 2018). In such settings, Axon-seq provides an additional layer of transcriptome-wide data, facilitating identification of the transcriptional networks underlying pathological changes in human motor axons. In our current system we have studied growing axons prior to forming neuromuscular junctions. Stage-specific changes in the retinal ganglion cell axonal mRNA repertoire were identified between growing, pruning and mature axons in the mouse brain (Shigeoka et al., 2016). It is to be expected that also the motor axon transcriptome will be influenced by the formation and maturity of a neuromuscular synapse. Applying Axon-seq to a microfluidic system containing iPSC-derived motor neurons and muscle could be used to investigate such processes in the context of the human NMJ and it’s instability in ALS.

## EXPERIMENTAL PROCEDURES

### Differentiation of mouse embryonic stem cells into motor neurons

mESCs were expanded on tissue culture dishes coated with 0.1% gelatin (Sigma-Aldrich) in ES media for two days: Knock-Out DMEM with 15% Knock-Out serum, 2mM L-glutamine, 100nM non-essential amino acids, 100 U/ml each of penicillin and streptomycin, 80nM β-mercaptoethanol (all Thermo Fisher) and 10^6^ U/ml leukaemia inhibitory factor (Merck-Millipore). Then, ES were dissociated with TrypLE Express (Thermo Fisher) and purified for 30 minutes on gelatin-coated plates. Embryoid body formation was conducted for 2 days by culturing mESCs in suspension (4×10^5^ cells/ml media) in neural differentiation media: 1:1 mix of Neurobasal and DMEM/F12 (Thermo Fisher) supplemented with 1× B27 (Thermo Fisher) 2mM L-glutamine, 100nM β-mercaptoethanol and 100 U/mL each of penicillin and streptomycin. During the initial day of embryoid body formation the media was kept in constant swirling motion at 30 rpm to ensure that uniformly sized spheres were formed. Embryoid bodies were subsequently patterned by the addition of 500nM of smoothened agonist (SAG, R&D) and 100nM retinoic acid (RA, Sigma-Aldrich) to the neural differentiation media for four days (Nichterwitz et al., 2016). Embryoid bodies containing motor neurons were dissociated for 20 minutes using TrypLE Express (Thermo Fisher), filtered through a 40 μm cell strainer (Fisher Scientific) and spun down for 5 minutes at 1,200 rpm for cell plating into the microfluidic devices.

### Culturing of mouse motor neurons in microfluidic devices

Microfluidic devices (SND150, Xona microfluidics) were attached to sterilized round cover glasses (28mm diameter, Menzel-Gläser) by drying these extensively and applying gentle force to seal them on the glass coverslips. Both compartments were then coated overnight with 15 μg/ml poly-L-ornithine (Sigma-Aldrich). The secondary coating consisted of a mix of 2 μg/ml of fibronectin (Sigma-Aldrich) and 10 μg/ml of laminin (Sigma-Aldrich).

Motor neurons were resuspended at 100,000 cells/μl and loaded into the microchannel in a 5μl droplet. At this point, neural differentiation media was supplemented with 10 ng/ml of GDNF, BDNF, CNTF and NT3, 200μM of ascorbic acid (Sigma-Aldrich), 100nM of RA and 2μM of 5-Fluoro-2′-deoxyuridine (5FdU, Sigma-Aldrich). Cells were allowed at least 30 minutes to attach in their compartment, after which media was added into the adjacent wells. The axonal compartment was filled with similar media, but instead containing 50 ng/ml of GDNF and BDNF to facilitate axonal recruitment. Volumes in the wells were adapted to ensure a flow across each chamber to gradually provide media, and to establish a flow and thus trophic factor gradient from the axonal compartment to the somatic compartment. Media in the devices was changed daily.

Axonal recruitment was stopped after 2 days. The concentrations of GDNF and BDNF in the axonal compartment were reduced to 5 ng/ml. In the motor neuron compartment, trophic factor concentrations were doubled from this point onwards, to 20 ng/ml. Additionally, RA was no longer included in the motor neuron media.

### Differentiation of human pluripotent stem cells into motor neurons

Human embryonic and induced pluripotent stem cells were maintained on Matrigel-coated culture dishes (Corning) and passaged with 5 μM Rock-inhibitor (Y-27632; Tocris). For differentiation into motor neurons, the protocol reported in (Guo et al., 2017) was used with minor adaptations. Briefly, cells were dissociated with TrypLE Express and resuspended as single cells in N2/B27 media to form embryoid bodies (DMEM-F12 supplemented with N2 in a 1:1 ratio with Neurobasal media supplemented with B27, all Thermo Fisher). For the first 2 days, this media was further supplemented with 5 μM Rock-inhibitor, 40 μM SB-431542, 200 nM LDN-193189 and 3 μM CHIR99021 (all Tocris) and 200 μM ascorbic acid (Sigma-Aldrich). As of the third day, the media was supplemented with 100 nM RA, 500 nM SAG and 200 μM ascorbic acid. On day 10 the EBs were dissociated for plating into microfluidic devices.

### Culturing of human motor neurons in microfluidic devices

For human motor neurons, devices with 450 μm grooves were used (SND450, Xona Microfluidics). The procedures were largely similar to culturing mouse motor neurons in microfluidic devices, with the following exceptions. Human motor neurons were plated at a lower density of 30.000 cells/μl (150.000 cells/device). After dissociation, cells were plated in devices in B27 media supplemented with 5 μM Rock-inhibitor, 200 μM ascorbic acid, 10 μM DAPT. 10 ng/ml of BDNF and GDNF were added to the somatodendritic compartment, while 50 ng/ml of BDNF and GDNF were added to the axonal compartment for recruitment of motor axons. After 3 days in devices, DAPT was excluded from the media and BDNF and GDNF levels were reduced to 10 ng/ml in the axonal compartment.

### Harvesting, library preparation and sequencing

Cultures were harvested after one week (mouse) or two weeks (human) in the microfluidic devices using 2% triton X-100 (in water) and 1.5U of RNAse inhibitor (Takara). Lysis solution (75 ml) was flushed through the compartments, collected and snap-frozen on dry ice. Prior to library preparation, the volume of axonal samples was reduced from ~75μl to 12-15μl using a concentrator (Eppendorf) and 10μl of the lysate used for reverse transcription. 5μl of the lysates from somatodendritic samples were directly used for the RT reaction, carried out in a final volume of 10 μl. Further steps were carried out as described (Nichterwitz et al., 2016) and samples sequenced on Illumina HiSeq2000 and −2500 platforms.

### RNA-seq data processing and analysis

Mouse sequencing reads were mapped to the mm10 mouse reference genome and the human SOD1 gene locus (hg38/GRCh38) with HISAT version 2.0.3 (Kim et al., 2015), while the hg38 reference genome was used for human samples. Aligned reads were extracted and assigned using the GenomicAlignments package (version 1.8.4, Lawrence et al., 2013) in R, with the function summarizeOverlaps, mode set to ‘union’. Normalized RPKM values were calculated using gene exon lengths from the respective reference genomes. Basic sequencing quality control was performed. Exclusion criteria were: < 300,000 (axons) or <500,000 (somas) uniquely mapped reads, Spearman correlation to other samples < 0.4 or < 2,500 genes expressed at RPKM > 0.1. To ensure that high-quality axonal samples were not biologically cross-contaminated with material from cell somas, during the cell seeding process or the lysis, we conducted Spearman correlation, unsupervised hierarchical clustering and PCA of all samples that passed initial QC. Axonal samples that clustered with soma samples in these analyses were removed. For the control Hb9::GFP mESC line, MN soma samples (N=13) and axon samples (N=8) were derived from a total of six independent experiments. For the SOD1^G93A^-overexpressing mESC line, MN soma samples (N=10) and axon samples (N=4) were derived from a total of six independent experiments. Human MN soma samples (N=7) and axon samples (N=6) were derived from two iPSC lines. The human single MN analysis was based on an N=39.

### RNAscope on mouse motor neurons

RNAscope (Wang et al., 2012) was used to verify the expression of Cox6a1 and Ybx1 based on the sequencing data. In brief, neural progenitors or motor neuron cultures, grown on glass coverslips or in microfluidic devices, were fixed with fresh PFA (4% in PBS) for 30min at 4°C. The RNAscope multiplex fluorescent kit v1 was used (Cat. No. 320850) according to manufacturer recommendations. To evaluate the procedure in neural progenitor and motor neurons cultures we tested the triple negative (Cat. 320871) and positive control probes (Polr2a-C1, Ppib-C2 and Ubc-C3) provided with the fluorescent kit in addition to a probe targeting poly-A tails of mRNAs (Cat. 318631-C2, 1 to 100 dilution). To easily visualize the motor neurons, we combined the RNAscope procedure with immunofluorescence. After the hybridization with the last probe (Amp 4-FL) samples were washed and stained with an anti-GFP antibody (see Supplemental Table S1).

Microfluidic devices were stained with Cox6a1 (Cat. 519781-C2), Ybx1 (548711-C1) or Nrp1 (471621-C1) probes and the anti-GFP antibody. Representative images were taken with a Zeiss LSM800 confocal microscope at the Biomedicum Imaging Core Facility, Karolinska Institutet.

## ACCESSION NUMBERS

The original RNA sequencing data from this study are available at the NCBI Gene Expression Omnibus (GEO, http://www.ncbi.nlm.nih.gov/geo/) under accession number GSE121069.

## SUPPLEMENTAL INFORMATION

Supplementary Data and Experimental Procedures are available online.

## ACKNOWLEDGEMENTS

We would like to thank Mattias Karlen for his excellent work in creating the schematic for figure 1. We would like to thank Dr Sebastian Thams for generously providing the mESC lines and professor Kevin Eggan for generously providing one human iPSC line and the human Hb9::GFP ESC line. This work was supported by grants from the Swedish Research Council [2016-02112]; EU Joint Programme for Neurodegenerative Disease (JPND) [5292014-7500]; the Strategic Research Programme in Neuroscience; Karolinska Institutet; Birgit Backmark’s Donation to ALS Research at Karolinska Institutet in memory of Nils and Hans Backmark; Åhlén-stiftelsen (mA1/h17); Ulla-Carin Lindquists stiftelse för ALS forskning; and Magnus Bergvalls stiftelse [2015-00783, 2016-01531] to E.H. J.A.B. is supported by a postdoctoral fellowship from the Swedish Society for Medical Research (SSMF) and N.K. is supported by a postdoctoral fellowship from the Swedish Brain Foundation. Funding for open access charge: Swedish Research Council [2016-02112] to E.H.

## AUTHOR CONTRIBUTIONS

E.H. conceived the project. J.N., J.A.B and E.H. designed experiments. J.N., J.A.B and R.H. acquired data. J.N., J.A.B, R.H., N.K. and E.H. analyzed data. E.H. supervised the project. J.N. and E.H. wrote the manuscript with the help of J.A.B. and N.K. All authors edited and gave critical input on the manuscript.

## References

Allodi, I., and Hedlund, E. (2014). Directed midbrain and spinal cord neurogenesis from pluripotent stem cells to model development and disease in a dish. Front Neurosci 8, 109.

Amiri, M., and Hollenbeck, P.J. (2008). Mitochondrial biogenesis in the axons of vertebrate peripheral neurons. Dev Neurobiol 68, 1348–1361.

Ben-Yaakov, K., Dagan, S.Y., Segal-Ruder, Y., Shalem, O., Vuppalanchi, D., Willis, D.E., Yudin, D., Rishal, I., Rother, F., Bader, M., et al. (2012). Axonal transcription factors signal retrogradely in lesioned peripheral nerve. EMBO J 31, 1350–1363.

Besse, F., and Ephrussi, A. (2008). Translational control of localized mRNAs: restricting protein synthesis in space and time. Nat Rev Mol Cell Biol 9, 971–980.

Bicker, S., Khudayberdiev, S., Weiss, K., Zocher, K., Baumeister, S., and Schratt, G. (2013). The DEAH-box helicase DHX36 mediates dendritic localization of the neuronal precursor-microRNA-134. Genes Dev 27, 991–996.

Bigler, R.L., Kamande, J.W., Dumitru, R., Niedringhaus, M., and Taylor, A.M. (2017). Messenger RNAs localized to distal projections of human stem cell derived neurons. Sci Rep 611.

Bizzoca, A., Corsi, P., and Gennarini, G. (2009). The mouse F3/contactin glycoprotein: structural features, functional properties and developmental significance of its regulated expression. Cell adhesion & migration 3, 53–63.

Boyden, S. (1962). The chemotactic effect of mixtures of antibody and antigen on polymorphonuclear leucocytes. J Exp Med 115, 453–466.

Briese, M., Saal, L., Appenzeller, S., Moradi, M., Baluapuri, A., and Sendtner, M. (2016). Whole transcriptome profiling reveals the RNA content of motor axons. Nucleic Acids Res 44, e33.

Campbell, D.S., and Holt, C.E. (2001). Chemotropic responses of retinal growth cones mediated by rapid local protein synthesis and degradation. Neuron 32, 1013–1026.

Campenot, R.B. (1977). Local control of neurite development by nerve growth factor. Proc Natl Acad Sci U S A 74, 4516–4519.

Chauvet, S., Cohen, S., Yoshida, Y., Fekrane, L., Livet, J., Gayet, O., Segu, L., Buhot, M.C., Jessell, T.M., Henderson, C.E., et al. (2007). Gating of Sema3E/PlexinD1 signaling by neuropilin-1 switches axonal repulsion to attraction during brain development. Neuron 56, 807–822.

Chhangani, D., Endo, F., Amanullah, A., Upadhyay, A., Watanabe, S., Mishra, R., Yamanaka, K., and Mishra, A. (2016). Mahogunin ring finger 1 confers cytoprotection against mutant SOD1 aggresomes and is defective in an ALS mouse model. Neurobiol Dis 86, 16–28.

Cifuentes-Diaz, C., Nicole, S., Velasco, M.E., Borra-Cebrian, C., Panozzo, C., Frugier, T., Millet, G., Roblot, N., Joshi, V., and Melki, J. (2002). Neurofilament accumulation at the motor endplate and lack of axonal sprouting in a spinal muscular atrophy mouse model. Hum Mol Genet 11, 1439–1447.

Comley, L.H., Nijssen, J., Frost-Nylen, J., and Hedlund, E. (2016). Cross-disease comparison of amyotrophic lateral sclerosis and spinal muscular atrophy reveals conservation of selective vulnerability but differential neuromuscular junction pathology. J Comp Neurol 524, 1424–1442.

Cox, L.J., Hengst, U., Gurskaya, N.G., Lukyanov, K.A., and Jaffrey, S.R. (2008). Intra-axonal translation and retrograde trafficking of CREB promotes neuronal survival. Nat Cell Biol 10, 149–159.

Dasen, J.S., De Camilli, A., Wang, B., Tucker, P.W., and Jessell, T.M. (2008). Hox repertoires for motor neuron diversity and connectivity gated by a single accessory factor, FoxP1. Cell 134, 304–316.

DeJesus-Hernandez, M., Mackenzie, I.R., Boeve, B.F., Boxer, A.L., Baker, M., Rutherford, N.J., Nicholson, A.M., Finch, N.A., Flynn, H., and Adamson, J. (2011). Expanded GGGGCC hexanucleotide repeat in noncoding region of C9ORF72 causes chromosome 9p-linked FTD and ALS. Neuron 72, 245–256.

Du, J., Tran, T., Fu, C., and Sretavan, D.W. (2007). Upregulation of EphB2 and ephrin-B2 at the optic nerve head of DBA/2J glaucomatous mice coincides with axon loss. Invest Ophthalmol Vis Sci 48, 5567–5581.

Fischer, L.R., Culver, D.G., Tennant, P., Davis, A.A., Wang, M., Castellano-Sanchez, A., Khan, J., Polak, M.A., and Glass, J.D. (2004). Amyotrophic lateral sclerosis is a distal axonopathy: evidence in mice and man. Exp Neurol 185, 232–240.

Garner, C.C., Tucker, R.P., and Matus, A. (1988). Selective localization of messenger RNA for cytoskeletal protein MAP2 in dendrites. Nature 336, 674–677.

Groen, E.J., Fumoto, K., Blokhuis, A.M., Engelen-Lee, J., Zhou, Y., van den Heuvel, D.M., Koppers, M., van Diggelen, F., van Heest, J., Demmers, J.A., et al. (2013). ALS-associated mutations in FUS disrupt the axonal distribution and function of SMN. Hum Mol Genet 22, 3690–3704.

Gumy, L.F., Yeo, G.S., Tung, Y.C., Zivraj, K.H., Willis, D., Coppola, G., Lam, B.Y., Twiss, J.L., Holt, C.E., and Fawcett, J.W. (2011). Transcriptome analysis of embryonic and adult sensory axons reveals changes in mRNA repertoire localization. RNA 17, 85–98.

Guo, W., Naujock, M., Fumagalli, L., Vandoorne, T., Baatsen, P., Boon, R., Ordovas, L., Patel, A., Welters, M., Vanwelden, T., et al. (2017). HDAC6 inhibition reverses axonal transport defects in motor neurons derived from FUS-ALS patients. Nature communications 861.

Hanz, S., Perlson, E., Willis, D., Zheng, J.Q., Massarwa, R., Huerta, J.J., Koltzenburg, M., Kohler, M., van-Minnen, J., Twiss, J.L., et al. (2003). Axoplasmic importins enable retrograde injury signaling in lesioned nerve. Neuron 40, 1095–1104.

Holt, C.E., and Schuman, E.M. (2013). The central dogma decentralized: new perspectives on RNA function and local translation in neurons. Neuron 80, 648–657.

Hornberg, H., Wollerton-van Horck, F., Maurus, D., Zwart, M., Svoboda, H., Harris, W.A., and Holt, C.E. (2013). RNA-binding protein Hermes/RBPMS inversely affects synapse density and axon arbor formation in retinal ganglion cells in vivo. J Neurosci 33, 1038410395.

Huettl, R.E., Soellner, H., Bianchi, E., Novitch, B.G., and Huber, A.B. (2011). Npn-1 contributes to axon-axon interactions that differentially control sensory and motor innervation of the limb. PLoS Biol 9, e1001020.

Ji, S.J., and Jaffrey, S.R. (2012). Intra-axonal translation of SMAD1/5/8 mediates retrograde regulation of trigeminal ganglia subtype specification. Neuron 74, 95–107.

Kadkhodaei, B., Alvarsson, A., Schintu, N., Ramskold, D., Volakakis, N., Joodmardi, E., Yoshitake, T., Kehr, J., Decressac, M., Bjorklund, A., et al. (2013). Transcription factor Nurr1 maintains fiber integrity and nuclear-encoded mitochondrial gene expression in dopamine neurons. Proc Natl Acad Sci U S A 110, 2360–2365.

Kalmar, B., Innes, A., Wanisch, K., Kolaszynska, A.K., Pandraud, A., Kelly, G., Abramov, A.Y., Reilly, M.M., Schiavo, G., and Greensmith, L. (2017). Mitochondrial deficits and abnormal mitochondrial retrograde axonal transport play a role in the pathogenesis of mutant Hsp27-induced Charcot Marie Tooth Disease. Hum Mol Genet 26, 3313–3326.

Kar, A.N., Vargas, J.N.S., Chen, C.Y., Kowalak, J.A., Gioio, A.E., and Kaplan, B.B. (2017). Molecular determinants of cytochrome C oxidase IV mRNA axonal trafficking. Mol Cell Neurosci 80, 32–43.

Kenna, K.P., van Doormaal, P.T., Dekker, A.M., Ticozzi, N., Kenna, B.J., Diekstra, F.P., van Rheenen, W., van Eijk, K.R., Jones, A.R., Keagle, P., et al. (2016). NEK1 variants confer susceptibility to amyotrophic lateral sclerosis. Nat Genet 48, 1037–1042.

Kim, D., Langmead, B., and Salzberg, S.L. (2015). HISAT: a fast spliced aligner with low memory requirements. Nat Methods 12, 357–360.

Kiskinis, E., Sandoe, J., Williams, L.A., Boulting, G.L., Moccia, R., Wainger, B.J., Han, S., Peng, T., Thams, S., Mikkilineni, S., et al. (2014). Pathways disrupted in human ALS motor neurons identified through genetic correction of mutant SOD1. Cell stem cell 14, 781–795.

Kwiatkowski, T.J., Bosco, D., Leclerc, A., Tamrazian, E., Vanderburg, C., Russ, C., Davis, A., Gilchrist, J., Kasarskis, E., and Munsat, T. (2009). Mutations in the FUS/TLS gene on chromosome 16 cause familial amyotrophic lateral sclerosis. Science 323, 1205–1208.

Lasek, R.J., Garner, J.A., and Brady, S.T. (1984). Axonal transport of the cytoplasmic matrix. J Cell Biol 99, 212s–221s.

Lee, S.K., and Hollenbeck, P.J. (2003). Organization and translation of mRNA in sympathetic axons. J Cell Sci 116, 4467–4478.

Lin, Y.C., and Koleske, A.J. (2010). Mechanisms of synapse and dendrite maintenance and their disruption in psychiatric and neurodegenerative disorders. Annu Rev Neurosci 33, 349–378.

Litman, P., Barg, J., Rindzoonski, L., and Ginzburg, I. (1993). Subcellular localization of tau mRNA in differentiating neuronal cell culture: implications for neuronal polarity. Neuron 10, 627–638.

Lyabin, D.N., Doronin, A.N., Eliseeva, I.A., Guens, G.P., Kulakovskiy, I.V., and Ovchinnikov, L.P. (2014a). Alternative forms of Y-box binding protein 1 and YB-1 mRNA. PLoS One 9, e104513.

Lyabin, D.N., Eliseeva, I.A., and Ovchinnikov, L.P. (2014b). YB-1 protein: functions and regulation. Wiley interdisciplinary reviews RNA 5, 95–110.

McIntyre, J.C., Titlow, W.B., and McClintock, T.S. (2010). Axon growth and guidance genes identify nascent, immature, and mature olfactory sensory neurons. J Neurosci Res 88, 3243–3256.

Mills, R., Taylor-Weiner, H., Correia, J.C., Agudelo, L.Z., Allodi, I., Kolonelou, C., Martinez-Redondo, V., Ferreira, D.M.S., Nichterwitz, S., Comley, L.H., et al. (2018). Neurturin is a PGC-1alpha1-controlled myokine that promotes motor neuron recruitment and neuromuscular junction formation. Mol Metab 7, 12–22.

Minis, A., Dahary, D., Manor, O., Leshkowitz, D., Pilpel, Y., and Yaron, A. (2014). Subcellular transcriptomics-dissection of the mRNA composition in the axonal compartment of sensory neurons. Dev Neurobiol 74, 365–381.

Naumann, M., Pal, A., Goswami, A., Lojewski, X., Japtok, J., Vehlow, A., Naujock, M., Gunther, R., Jin, M., Stanslowsky, N., et al. (2018). Impaired DNA damage response signaling by FUS-NLS mutations leads to neurodegeneration and FUS aggregate formation. Nature communications 9, 335.

Nichterwitz, S., Chen, G., Aguila Benitez, J., Yilmaz, M., Storvall, H., Cao, M., Sandberg, R., Deng, Q., and Hedlund, E. (2016). Laser capture microscopy coupled with Smart-seq2 for precise spatial transcriptomic profiling. Nat Commun 7, 12139.

Picelli, S., Bjorklund, A.K., Faridani, O.R., Sagasser, S., Winberg, G., and Sandberg, R. (2013). Smart-seq2 for sensitive full-length transcriptome profiling in single cells. Nature methods 10, 1096–1098.

Renton, A.E., Majounie, E., Waite, A., Simón-Sánchez, J., Rollinson, S., Gibbs, J.R., Schymick, J.C., Laaksovirta, H., Van Swieten, J.C., and Myllykangas, L. (2011). A hexanucleotide repeat expansion in C9ORF72 is the cause of chromosome 9p21-linked ALS-FTD. Neuron 72, 257–268.

Rosen, D.R., Siddique, T., Patterson, D., Figlewicz, D.A., Sapp, P., Hentati, A., Donaldson, D., Goto, J., O’Regan, J.P., and Deng, H.-X. (1993). Mutations in Cu/Zn superoxide dismutase gene are associated with familial amyotrophic lateral sclerosis.

Rotem, N., Magen, I., Ionescu, A., Gershoni-Emek, N., Altman, T., Costa, C.J., Gradus, T., Pasmanik-Chor, M., Willis, D.E., Ben-Dov, I.Z., et al. (2017). ALS Along the Axons - Expression of Coding and Noncoding RNA Differs in Axons of ALS models. Sci Rep 7, 44500.

Saal, L., Briese, M., Kneitz, S., Glinka, M., and Sendtner, M. (2014). Subcellular transcriptome alterations in a cell culture model of spinal muscular atrophy point to widespread defects in axonal growth and presynaptic differentiation. RNA 20, 1789–1802.

Sgado, P., Alberi, L., Gherbassi, D., Galasso, S.L., Ramakers, G.M., Alavian, K.N., Smidt, M.P., Dyck, R.H., and Simon, H.H. (2006). Slow progressive degeneration of nigral dopaminergic neurons in postnatal Engrailed mutant mice. Proc Natl Acad Sci U S A 103, 15242–15247.

Sharma, S., Grintsevich, E.E., Hsueh, C., Reisler, E., and Gimzewski, J.K. (2012). Molecular cooperativity of drebrin1-300 binding and structural remodeling of F-actin. Biophys J 103, 275–283.

Shigeoka, T., Jung, H., Jung, J., Turner-Bridger, B., Ohk, J., Lin, J.Q., Amieux, P.S., and Holt, C.E. (2016). Dynamic Axonal Translation in Developing and Mature Visual Circuits. Cell 166, 181–192.

Sreedharan, J., Blair, I.P., Tripathi, V.B., Hu, X., Vance, C., Rogelj, B., Ackerley, S., Durnall, J.C., Williams, K.L., and Buratti, E. (2008). TDP-43 mutations in familial and sporadic amyotrophic lateral sclerosis. Science 319, 1668–1672.

Sugiyama, S., Di Nardo, A.A., Aizawa, S., Matsuo, I., Volovitch, M., Prochiantz, A., and Hensch, T.K. (2008). Experience-dependent transfer of Otx2 homeoprotein into the visual cortex activates postnatal plasticity. Cell 134, 508–520.

Swinnen, B., and Robberecht, W. (2014). The phenotypic variability of amyotrophic lateral sclerosis. Nat Rev Neurol 10, 661–670.

Taylor, A.M., Berchtold, N.C., Perreau, V.M., Tu, C.H., Li Jeon, N., and Cotman, C.W. (2009). Axonal mRNA in uninjured and regenerating cortical mammalian axons. J Neurosci 29, 4697–4707.

Taylor, A.M., Blurton-Jones, M., Rhee, S.W., Cribbs, D.H., Cotman, C.W., and Jeon, N.L. (2005). A microfluidic culture platform for CNS axonal injury, regeneration and transport. Nature methods 2, 599–605.

Telley, L., Cadilhac, C., Cioni, J.M., Saywell, V., Jahannault-Talignani, C., Huettl, R.E., Sarrailh-Faivre, C., Dayer, A., Huber, A.B., and Ango, F. (2016). Dual Function of NRP1 in Axon Guidance and Subcellular Target Recognition in Cerebellum. Neuron 91, 1276–1291.

Torero Ibad, R., Rheey, J., Mrejen, S., Forster, V., Picaud, S., Prochiantz, A., and Moya, K.L. (2011). Otx2 promotes the survival of damaged adult retinal ganglion cells and protects against excitotoxic loss of visual acuity in vivo. J Neurosci 31, 5495–5503.

Upadhyay, A., Amanullah, A., Chhangani, D., Mishra, R., Prasad, A., and Mishra, A. (2016). Mahogunin Ring Finger-1 (MGRN1), a Multifaceted Ubiquitin Ligase: Recent Unraveling of Neurobiological Mechanisms. Mol Neurobiol 53, 4484–4496.

Valdez, G., Tapia, J.C., Lichtman, J.W., Fox, M.A., and Sanes, J.R. (2012). Shared resistance to aging and ALS in neuromuscular junctions of specific muscles. PLoS One 7, e34640.

Vance, C., Rogelj, B., Hortobágyi, T., De Vos, K.J., Nishimura, A.L., Sreedharan, J., Hu, X., Smith, B., Ruddy, D., and Wright, P. (2009). Mutations in FUS, an RNA processing protein, cause familial amyotrophic lateral sclerosis type 6. Science 323, 1208–1211.

Venkova, K., Christov, A., Kamaluddin, Z., Kobalka, P., Siddiqui, S., and Hensley, K. (2014). Semaphorin 3A signaling through neuropilin-1 is an early trigger for distal axonopathy in the SOD1G93A mouse model of amyotrophic lateral sclerosis. J Neuropathol Exp Neurol 73, 702–713.

Wang, F., Flanagan, J., Su, N., Wang, L.C., Bui, S., Nielson, A., Wu, X., Vo, H.T., Ma, X.J., and Luo, Y. (2012). RNAscope: a novel in situ RNA analysis platform for formalin-fixed, paraffin-embedded tissues. The Journal of molecular diagnostics: JMD 14, 22–29.

Wichterle, H., Lieberam, I., Porter, J.A., and Jessell, T.M. (2002). Directed differentiation of embryonic stem cells into motor neurons. Cell 110, 385–397.

Wizenmann, A., Brunet, I., Lam, J., Sonnier, L., Beurdeley, M., Zarbalis, K., Weisenhorn-Vogt, D., Weinl, C., Dwivedy, A., Joliot, A., et al. (2009). Extracellular Engrailed participates in the topographic guidance of retinal axons in vivo. Neuron 64, 355–366.

Yoon, B.C., Jung, H., Dwivedy, A., O’Hare, C.M., Zivraj, K.H., and Holt, C.E. (2012). Local translation of extranuclear lamin B promotes axon maintenance. Cell 148, 752–764.

Zhang, Y., Chen, K., Sloan, S.A., Bennett, M.L., Scholze, A.R., O’Keeffe, S., Phatnani, H.P., Guarnieri, P., Caneda, C., Ruderisch, N., et al. (2014). An RNA-sequencing transcriptome and splicing database of glia, neurons, and vascular cells of the cerebral cortex. J Neurosci 34, 11929–11947.

Zhuang, B., Su, Y.S., and Sockanathan, S. (2009). FARP1 promotes the dendritic growth of spinal motor neuron subtypes through transmembrane Semaphorin6A and PlexinA4 signaling. Neuron 61, 359–372.

Zuko, A., Oguro-Ando, A., Post, H., Taggenbrock, R.L., van Dijk, R.E., Altelaar, A.F., Heck, A.J., Petrenko, A.G., van der Zwaag, B., Shimoda, Y., et al. (2016). Association of Cell Adhesion Molecules Contactin-6 and Latrophilin-1 Regulates Neuronal Apoptosis. Front Mol Neurosci 9, 143

